# ‘Fishing’ for Mitochondrial DNA in The Egyptian Sacred Ibis Mummies

**DOI:** 10.1101/473454

**Authors:** Sally Wasef, Leon Huynen, Craig Donald Millar, Sankar Subramanian, Salima Ikram, Barbra Holland, Eske Willerslev, David Martin Lambert

## Abstract

The long-term preservation of DNA requires a number of optimal conditions, including consistent exposure to cool, dry, and dark environments. As a result, the successful recovery of ancient DNA from material from warmer climates such as those in Egypt has often been met with scepticism. Egypt has an abundance of ancient mummified animals and humans, whose genetic analyses would offer important insights into ancient cultural practices. To date, the retrieval of complete genomes from ancient Egyptian remains of humans or other animals has been largely unsuccessful. To test for the presence of even short DNA sequences in Egyptian material, we performed second-generation shotgun sequencing of DNA libraries constructed from ancient Sacred Ibis mummies. Since most of the resulting Illumina libraries were shown to contain extremely low levels (less than 0.06%) of endogenous mitochondrial DNA, we aimed to enrich these samples using targeted in-solution hybridisation methods. Using biotinylated RNA baits designed to Sacred Ibis complete mitochondrial sequences, we trialled a number of conditions and parameters and achieved up to 4705-fold enrichment. We also found that a combination of hybridisation temperature and the use of the polymerase KAPA HiFi significantly increased both the efficiency of targeted hybridisation and post-hybridisation amplification respectively. Furthermore, improved enrichment was accompanied with only minor increases in clonality. Our method enabled us to reconstruct the first complete mitochondrial genomes from ancient Egyptian sub-fossil material.

## Introduction

Ancient Egyptians mummified many kinds of animals for a range of purposes, these included their beloved pets, animals that were sacred representations of specific gods, and ‘votive offerings’ – animal gifts presented to the gods (Ikram 2015a). Many votive offerings were of the Sacred Ibis (*Threskiornis aethiopicus*). Several million Ibis mummies were offered to Thoth, the god of writing and wisdom (Ikram 2015a; 2015b). Sacred Ibis can no longer be found in Egypt, becoming extinct by the end of the 19^th^ century (Meinertzhagen 1930: 438)

Genetic analyses of mummified Egyptian animals have been notoriously difficult due to the warm and, in places, humid climate of Egypt, conditions which are generally detrimental to the survival of DNA (Gilbert et al. 2005).

Despite these difficulties, preliminary successes have been achieved with studies of both mummified crocodiles (Hekkala et al. 2011) and cat remains (Ottoni et al. 2017; Kurushima et al. 2012). Although Schueneman et al. (2017) have had some success with retrieving DNA, it should be noted that from more than 160 samples tested, only three of them successfully retrieved SNP data. Both studies using the Polymerase Chain Reaction successfully amplified and sequenced a small number of very short fragments of mitochondrial DNA (mtDNA). The recovered sequences allowed the identification of the species and the establishment of genetic relationships between ancient Egyptian and modern day species (Hekkala et al. 2011; Kurushima et al. 2012). However, subsequent work using second-generation DNA sequencing has been largely unsuccessful in obtaining significant coverage of ancient Egyptian animal nuclear or mitochondrial genomes (Khairat et al. 2013).

Ancient DNA (aDNA), existing in very low concentrations if at all in archaeological materials from harsh climates, is typically fragmented and damaged, and contains high levels of contamination (Knapp & Hofreiter 2010). Such samples require improved extraction (Dabney et al. 2013) and library construction protocols (Gansauge & Meyer 2013). When this study was carried out in 2014, the Gansauge and Meyer (2013) method was prevalent. Subsequently more work has been done using other methods, such as Gansauge et al. (2017) and Glocke et al. (2017). This approach, in conjunction with targeted hybridisation enrichment systems adapted for ancient DNA have been shown to result in the retrieval of orders of magnitude more sequence data, particularly from short, highly contaminated, and limited ancient DNA samples (Briggs et al. 2009; Fu et al. 2013; Maricic et al. 2010,).

Targeted enrichment typically uses biotinylated DNA or RNA baits complementary to the target DNA or RNA of interest (Gnirke et al. 2009). In a typical hybridisation reaction, biotinylated baits are hybridised to the desired DNA fragment, and are then captured by binding to streptavidin beads (Gnirke et al. 2009). The targeted sequences that have been hybridised to the DNA/RNA baits are then released by denaturation with hydroxide, and amplified and sequenced using a second-generation platform (Mamanova et al. 2010). The efficiency of the capture system is commonly expressed as x-fold enrichment of target DNA.

Unfortunately, for reasons unknown, enrichment rates for different samples using the same experimental conditions often differ, sometimes dramatically. Enrichment rates vary particularly when targeting mtDNA, where rates have been shown to vary from 22 to 2217 fold (Enk et al. 2013). Parameters that may influence enrichment efficiency include the enzymes and chemistry used, sequence similarity between the baits and target, depth of bait tiling, and hybridisation and washing temperatures (Avila-Arcos et al. 2011; Bodi et al. 2013, Li et al. 2013). Research by Li et al. (2013) showed that for MYcroarray’s MYbaits™ in-solution capture system, a gradual decrease in hybridisation temperature (touchdown) significantly improves capture efficiency (Gnirke et al. 2009). Pajimans et al. (2015) also showed the importance of hybridisation temperature when dealing with different sample types.

In this pilot study, we attempt to retrieve complete mitochondrial genomes from ancient mummified Sacred Ibis tissue. Previous studies with Egyptian mummified animal samples (see above) showed that the amount of endogenous DNA in tissues was extremely low, precluding significant analyses. In this work, we used targeted hybridization to extract endogenous Sacred Ibis mitochondrial DNA. We also varied a number of parameters using MYcroarray’s MyBaits™ in-solution capture RNA hybridisation baits to enhance the retrieval of selected DNAs from ancient Egyptian mummified Sacred Ibis bone, tissue, and feathers.

## Material and Methods

### Sacred Ibis samples

Bone, tissue and feather samples were collected for research purposes from the main Egyptian Sacred Ibis catacombs at Saqqara, Tuna El-Gebel, and Sohag (Fig. 1, Tab. 1) with permission from Egypt’s Ministry of State for Antiquities. Further Sacred Ibis samples were obtained from a number of museums (Table 1). The age of the mummified ibises were estimated from a subset of the museum samples ranged between c.450 and 250 cal BC (Wasef et al. 2015).

**Table 1:**
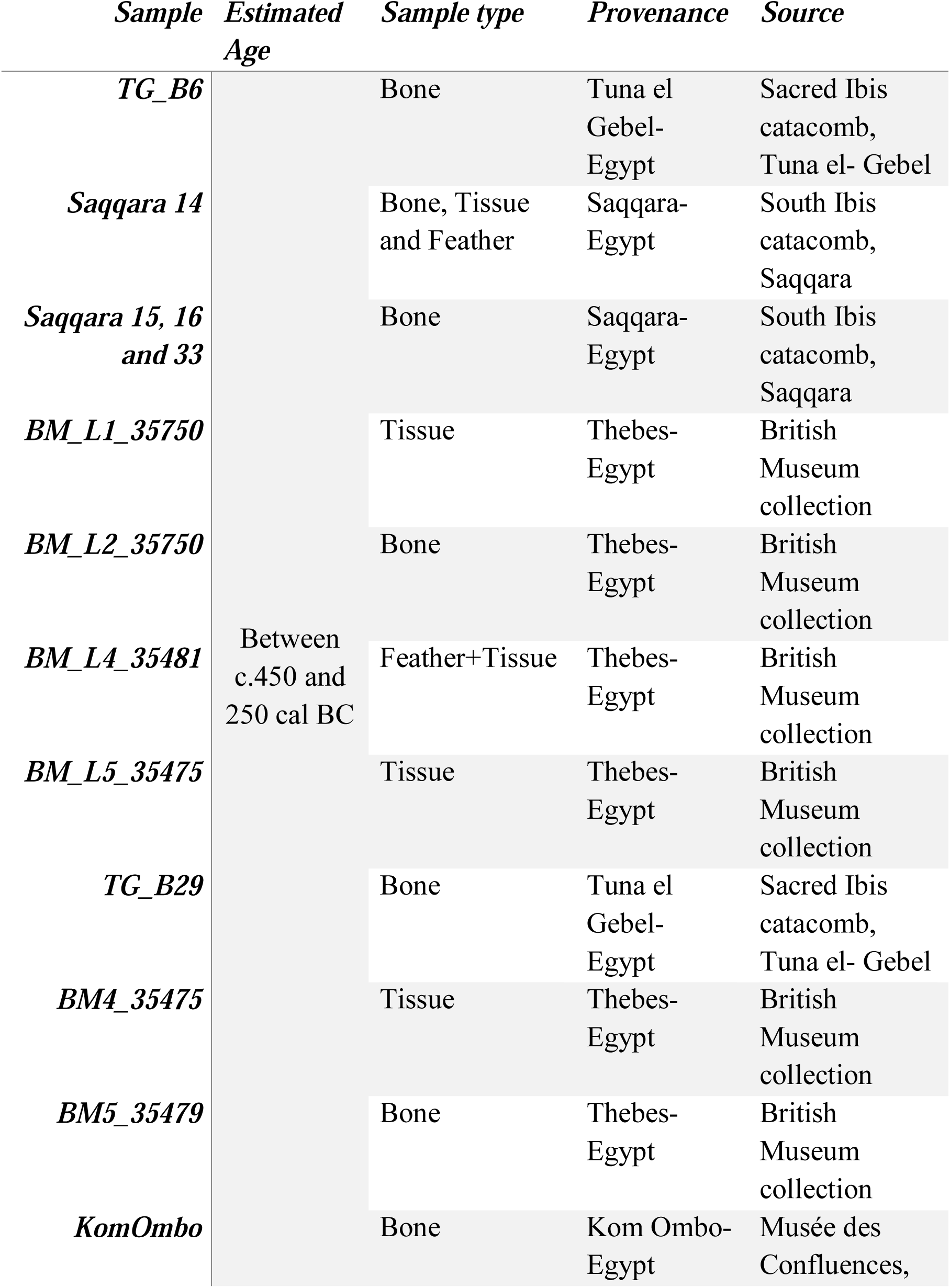
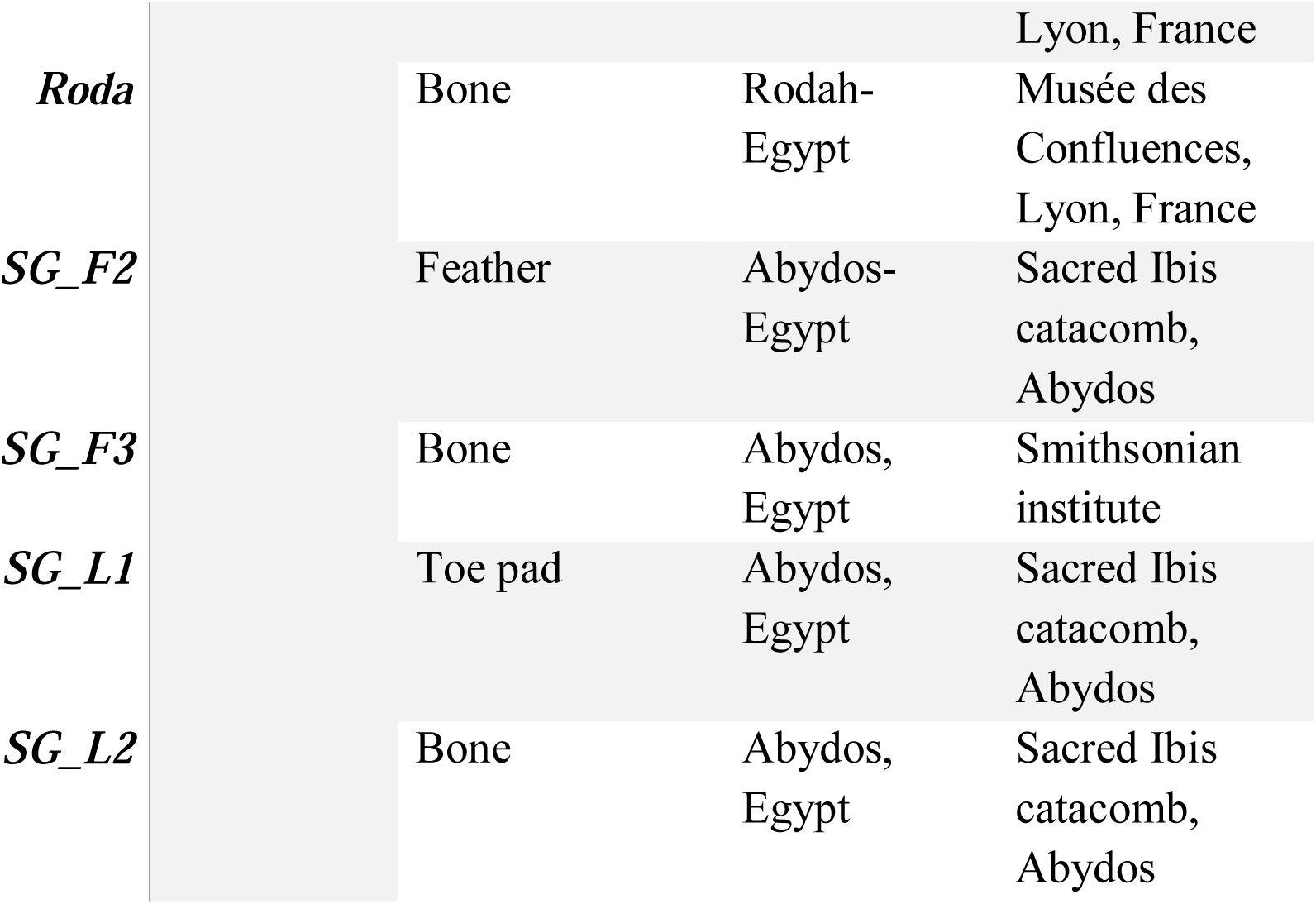
Sacred Ibis samples. Details of the location, tissue type and estimated ages of the ancient Sacred Ibis samples are shown.

**Fig. 1:**
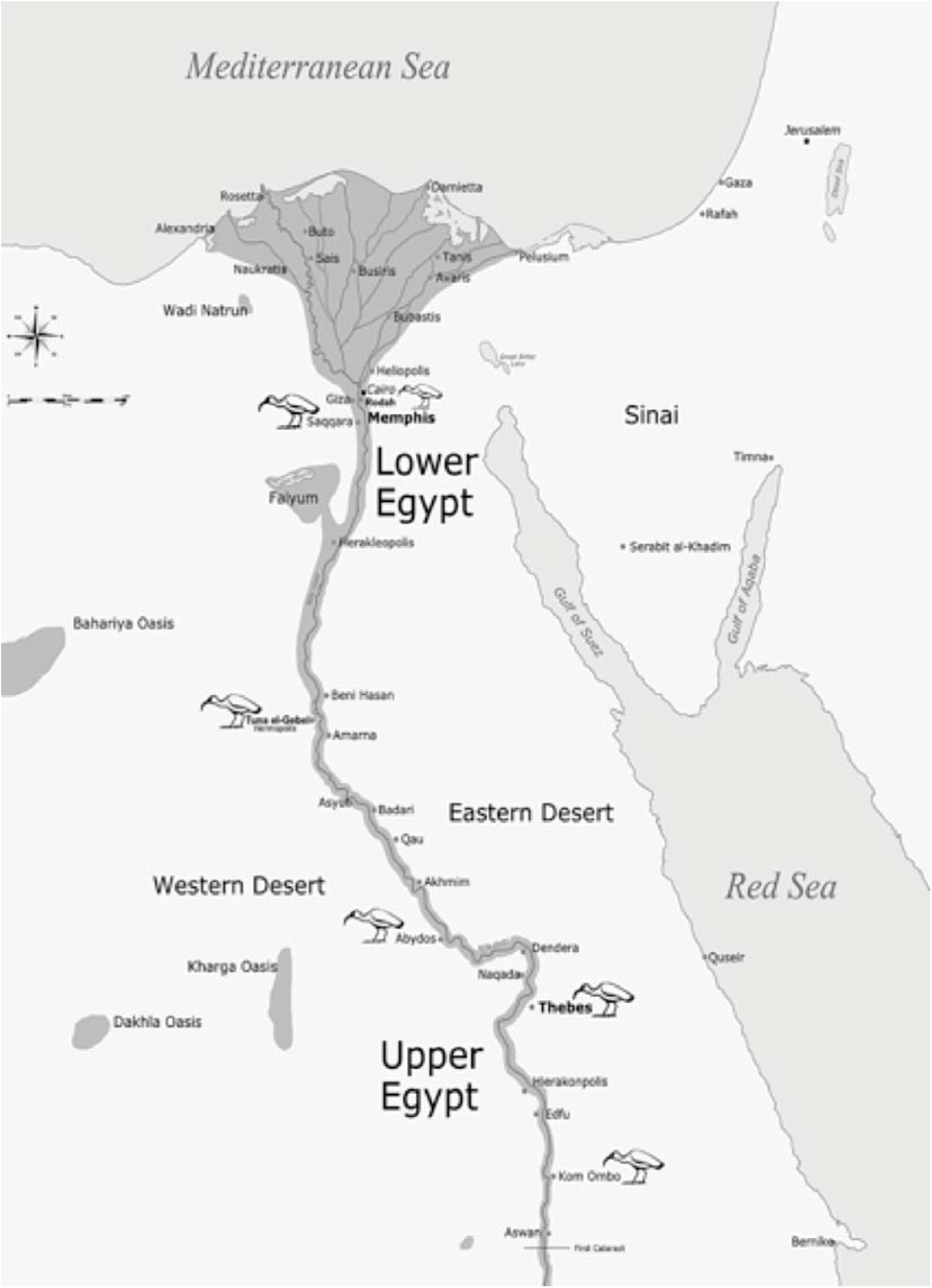
The location of archaeological (catacomb) sites in Egypt from which ancient Sacred Ibis mummified material was sampled. Map created by the authors.

### DNA extraction

Ancient Sacred Ibis samples were processed in accordance with requirements for handling ancient DNA as outlined by Knapp et al. (2012). For Sacred Ibis samples collected from Egyptian catacombs (Tab. 1) preliminary DNA extractions were carried out at an ancient DNA facility at Al Kasr Al Aini Medical School in Cairo, Egypt, while other Sacred Ibis ancient samples from international museums were extracted for DNA in the ancient DNA Laboratory facility at Griffith University, Brisbane, Australia. Ancient bone samples were initially treated with 10% bleach and then 80% alcohol to remove surface contamination. The outer layer was then removed and the remaining bone fragment was homogenized using a mortar and a pestle into fine powder. Approximately 30 mg of bone powder was digested in 600 μL of extraction buffer (0.45 M EDTA, 0.5% N-lauryl sarcosine, 1 mg/mL proteinase K and 20 uL of 1 M DTT) for 24hrs at 40 ^0^C with rotation. The residual bone powder was pelleted by centrifugation and the DNA in the supernatant was extracted several times with buffer-equilibrated phenol (pH 7.5) and then chloroform. Ten volumes of PB buffer (Qiagen) were then added to the extract (Dabney et al. 2013) and the DNA was purified using Qiagen DNeasy Blood & Tissue columns, as recommended by the manufacturer. Purified DNA was then eluted from the column using 70 μL of ultra-pure water.

In the case of ancient toe pads, soft tissue, and feathers samples, these were sliced with a scalpel to increase the surface area exposed to the extraction buffer. To those samples we added 200 uL of SET buffer, 40 uL of 10% SDS, 20 uL of 20 mg/ml Proteinase and 20 uL of 1 M DTT. Samples were incubated at 40 ^0^C, overnight with rotation, until completely digested. The extract was then purified following the same steps as for ancient bone. Final DNA concentrations and sizes were determined using a Qubit^®^ 2.0 Fluorometer and a Bioanalyzer 2100 (Agilent Technologies, Palo Alto, CA, US) respectively.

### Illumina sequencing library preparation

#### For ancient DNA

Ancient DNA Illumina sequencing libraries were constructed using a NEBNext^®^ DNA Library Preparation Kit with modifications as proposed by Meyer and Kircher (2010). Illumina Libraries prepared by this method from the ancient samples were used for both direct sequencing (shotgun sequencing) and target capture hybridisation reactions. A solution of 22 ul of aDNA was end repaired for 30 mins at 37°C then purified using a Qiagen Minelute column with 10 volumes of Qiagen PN or PB buffer and finally eluted in 18 uL of ultrapure water. 17 uL of the end repaired DNA was then ligated to blunt-end Illumina specific adapters (P5F, 5’-A*C*A*C*TCTTTCCCTACACGACGCTCTTCCG*A*T*C*T; P5+P7.R, 5’-A*G*A*T*CGGAA*G*A*G*C; P7F, 5’-G*T*G*A*CTGGAGTTCAGACGTGTGCTCTTCCG*A*T*C*T) in a 30 uL reaction and incubated for 25 mins at 20^0^C. Libraries were then purified using Qiagen Minelute columns. Adapter fill-in reactions were carried out with BstI polymerase in a final volume of 25 uL. The mix was incubated for 20 min at 65°C, before the BstI was heat inactivated at 80°C for 20 min. The ancient DNA Illumina libraries were then amplified in a 50 uL PCR reaction using 15 uL of the library template. Amplifications were carried out using either Phusion^®^ High-Fidelity PCR Master Mix in GC Buffer (NEB) for 20 to 22 cycles or KAPA HiFi Uracil+ polymerase Master Mix (KAPABiosystems) for 10 to 17 cycles. Amplified libraries were purified using 1 x Agencourt AMPure XP beads and then analysed using a Bioanalyzer 2100.

### Target capture hybridisation

A MYcroarray MyBaits™ in-solution capture system was used to recover complete mitochondrial genomes from ancient Sacred Ibis samples. Capture Baits to the complete Sacred Ibis mitochondrial genome were designed as 80- mer biotinylated RNAs with 5 base overlaps by MYcroarray. Approximately 100 – 500 ng of amplified Illumina library was denatured and then hybridised to adapter blocking primers. The blocked library was then hybridised to the single-stranded RNA baits at constant temperature. In the case of the ancient samples, we tested a range of hybridisation temperatures (45°C, 55°C, 57°C or 65°C). Incubation was carried out for 48hrs before being bound to magnetic streptavidin beads. The beads were washed with buffer heated to same temperature as that used for hybridisation to remove non-specifically bound DNAs. The captured DNA was eluted from the RNA baits with 100 mM NaOH and then neutralized with 1 M Tris-Cl pH 8.0. Finally, the DNA was purified using a Qiagen MinElute column after amplification.

### Illumina second generation sequencing

The sequencing process was exactly the same for either directly sequenced (shotgun) libraries or the amplified target enriched libraries: Indexed libraries were purified using 1 x Agencourt AMPure XP beads, then quantified and visualized using a Bioanalyzer 2100. Three to six libraries were then pooled together in equimolar amounts and sequenced using either an Illumina MiSeq sequencer at Griffith University as a single-end reads for 150 cycles, or were sequenced as single-end reads for 100 cycles using a single lane of an Illumina HiSeq2000 at the Danish National High-Throughput DNA Sequencing Facility in Copenhagen.

### NGS data processing

Sequence reads were initially analysed using the FASTX-Toolkit V0.0.13. Reads shorter than 25 bases and adapter sequences were removed and low quality bases were trimmed. Following these initial analyses, reads were aligned to the Sacred Ibis mitochondrial reference genome (GenBank accession number: NC 013146.1) using BWA V0.6.2-r126 (Li and Durbin, 2009) using bwa aln -f command. SAMtools was used to extract data, index, sort, and view output files (Li and Durbin, 2009). Qualimap (Okonechnikov, et al., 2015) was used to assess alignment quality. The presence of endogenous ancient DNA was determined by using mapDamage2.0 (Jonsson, et al., 2013) to measure nucleotide mis incorporations typical of ancient DNA. All mapped reads were used to determine mitogenome coverage and the consensus sequence.

## Results and Discussion

### Pre-capture Results

Using an aDNA extraction method (Dabney et al. 2013) and library building protocol specific for aDNA (Meyer and Kircher 2010), we reconstructed ancient second generation sequencing DNA libraries from various Sacred Ibis tissues. Initial direct (shotgun) sequencing of these libraries showed that the highest level of endogenous mitochondrial DNA (calculated as a percentage of unique sequences versus total number of reads) was recovered from ancient toe pad (x□ = 0.01%), followed by ancient feather and bone (x□ = 0.002%), and finally ancient soft tissue (x□= 0.0002%). The length of recovered DNA sequences varied significantly amongst the ancient samples, with DNA from bone averaging 63 ± 16 bp, soft tissue averaging 45.2 ± 11 bp, and single toe pads and feathers averaging 44.9 and 53.9 bp respectively. These results suggest that initial DNA degradation is likely to be the result of rapid endonuclear digestion followed by slow exonuclear ‘nibbling’, based on the difference in base pair length in the different tissues. Duplicates varied considerably from 3.03% to 89.26% with no significant differences noted between the various tissues. Biases in duplicate amplification also appear to be influenced by the polymerase used. Libraries amplified using KAPA HiFi Uracil+ have been shown to display clearer damage and fragmentation patterns characteristic of endogenous ancient DNA (Fig. 2) (Jonsson et al. 2013). As Phusion polymerase can have issues with the deaminated cytosines by blocking the correspondence of damages sites, KAPA was used to bypass the damage. Thus, we show that pre-capture amplification using KAPA HiFi Uracil+ polymerase resulted in the production of more unique sequences as well as more duplicates (Table 2), in comparison to when Phusion^®^ High-Fidelity polymerase was used.

**Table 2:**
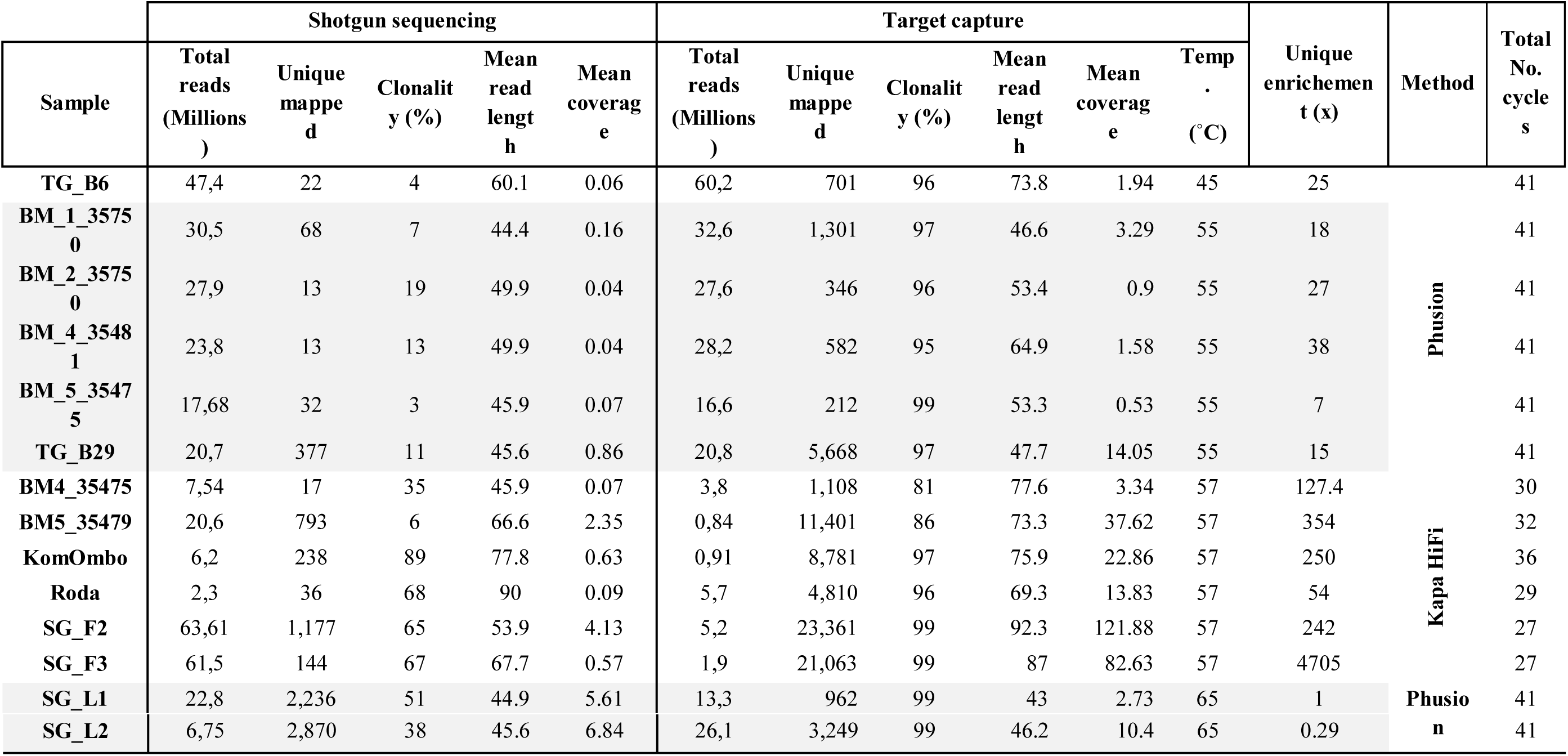
Description of data generated using various Ancient Sacred Ibis tissues showing: Unique mitochondrial sequences, mitogenome coverage and clonality for shotgun sequences; Unique mitochondrial sequences, mitogenome coverage, enrichment and clonality for captured sequences. ‘Total reads’ refers to the sequence data generated before alignment to the Sacred Ibis mitochondrial genome. ‘Unique’ refers to the fraction of mapped mitochondrial reads after the removal of clonal sequences. The enrichment rate (x fold) is calculated by dividing the % of unique endogenous mitochondrial sequences after capture by the total number of shotgun sequences.

**Fig. 2:**
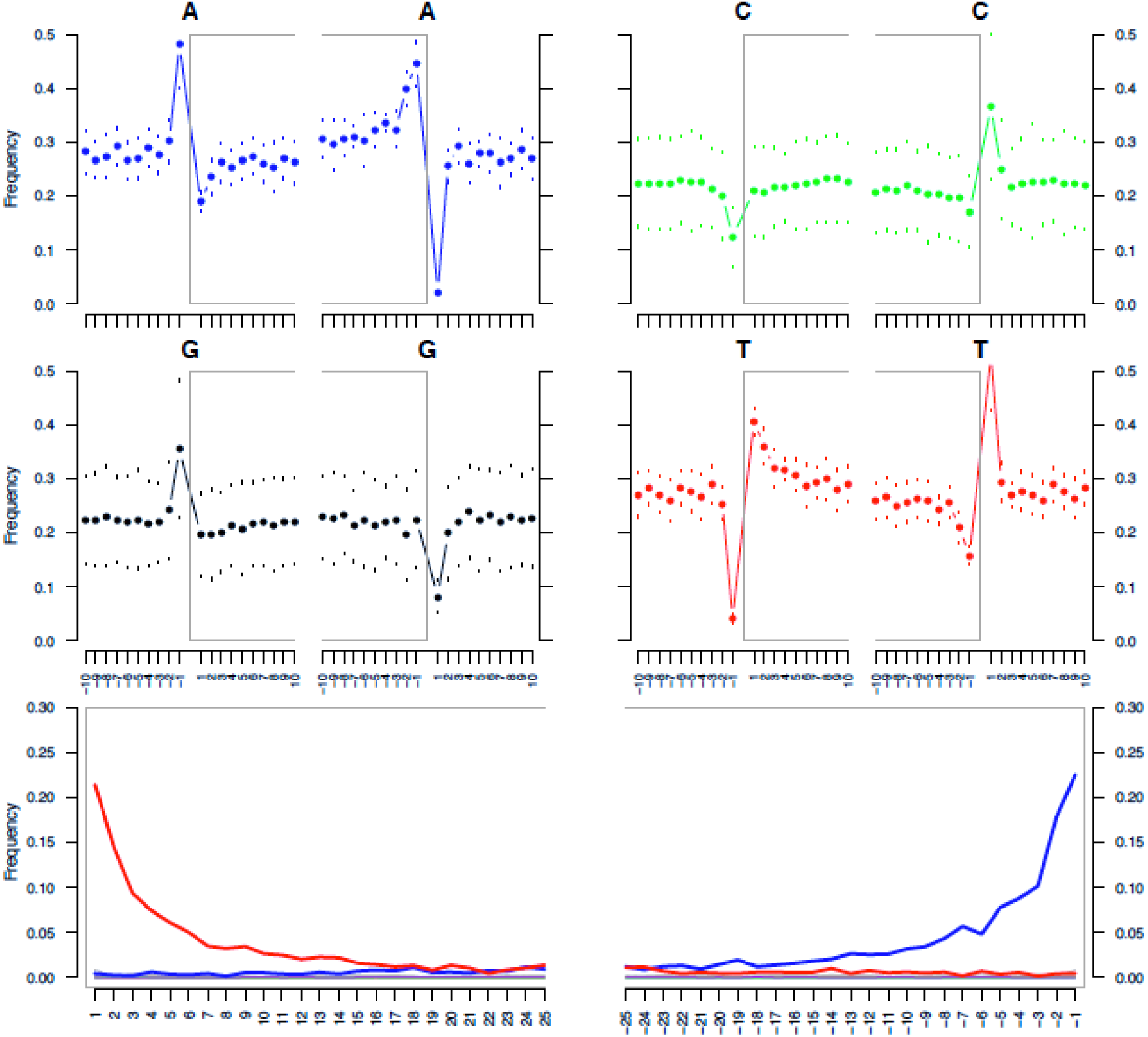
Nucleotide misincorporation patterns observed in the Sacred Ibis ancient library from Tuna El-Gebel. Damage is represented at the 3′ terminus by guanine to adenine substitutions (G>A; blue) and at the 5′ terminus by cytosine to thymine substitutions (C>T; red. The four upper mini-plots show the base frequency outside and in the read (the open grey box corresponds to the read). The bottom plots are the positions’ specific substitutions from the 5‴ (left) and the 3‴ end (right).

### Capture Results

Only samples that showed genuine ancient Ibis endogenous contents in the screening process using shotgun sequencing have been used for further enrichment by target capture baits. As a result of excluding other samples of low endogenous content and also the sequencing costs involved, we compared the simultaneous effects of altering a number of parameters (eg Phusion^®^ High-Fidelity polymerase or KAPA HiFi Uracil+ polymerase as well as different hybridisation temperatures) on the enrichment of endogenous DNA amongst the samples. The levels of endogenous DNA among the ancient samples before enrichment (using direct sequencing) was shown not to be statistically different (P=0.051), suggesting that there is no sample-specific bias, which means when directly sequencing different samples we would expect same results. We successfully retrieved fourteen complete ancient mitochondrial genomes from the mummified Sacred Ibis remains.

### Target Capture Enrichment Rates

Enrichment rates were calculated for each sample by comparing the computed percentage of the unique (non-clonal) sequences aligned to the Sacred Ibis mitochondrial reference genome pre and after capture hybridisation enrichment (Tab. 2). We found that regardless of the sample type used, the Taq polymerase in the library build method or the hybridisation temperature, an increase in the unique endogenous content of the captured libraries in comparison to the shotgun-sequenced counterpart (Tab. 2).

For ancient DNA a significant difference in enrichment efficiency was found to be dependent on the polymerase used. The polymerase KAPA HiFi Uracil+ enriched between 54 x - 4705 x requiring only 26 – 37 cycles, while enrichment using Phusion^®^ High-Fidelity PCR Master Mix in GC Buffer (NEB) resulted in only 0.3 – 38 x enrichment and required up to 41 cycles (see Tab. 2).

### Mean Read Length

A MYcroarray MyBaits^™^ in-solution capture system was used to enrich for Sacred Ibis mitochondrial DNA. To test for the efficiency of the bait length and tiling system to captured DNAs of different fragment sizes, and to measure the variation in mean read lengths after capture, we designed our biotinylated RNA baits to be 80-mer with a 5 base pair tiling depth. By comparing the insert size of the ancient pre-capture sequences to their equivalent post-capture sequences, we found a slight increase in the mean read length of the unique sequences for most samples (1.2 fold). Those results are consistent with previous observations (Carpenter et al. 2013; Enk et al. 2013 and 2014).

### Hybridisation Temperature

In addition to using a hybridisation temperature of 65°C, as recommended by the manufacturer, we also tested enrichment efficiencies at 45°C, 55°C, and 57°C. All post-capture washes were carried out at the same temperature as the hybridisation temperature. Enrichment rates were calculated for each sample by comparing the percentage of unique Sacred Ibis mitochondrial sequences pre- and post-enrichment (Tab. 2). In contrast to results published by Paijmans et al. (Paijmans et al. 2015), we show that the best enrichment temperature for ancient DNA was 57°C (Fig. 3), with enrichment rates between 54 x to 4705 x resulting in up to 121 x coverage of the Sacred Ibis mitogenome (Tab. 2). Although not tested here, further enrichment might be achieved using touchdown hybridisation, shown to be effective by Li et al. (2013).

**Fig. 3:**
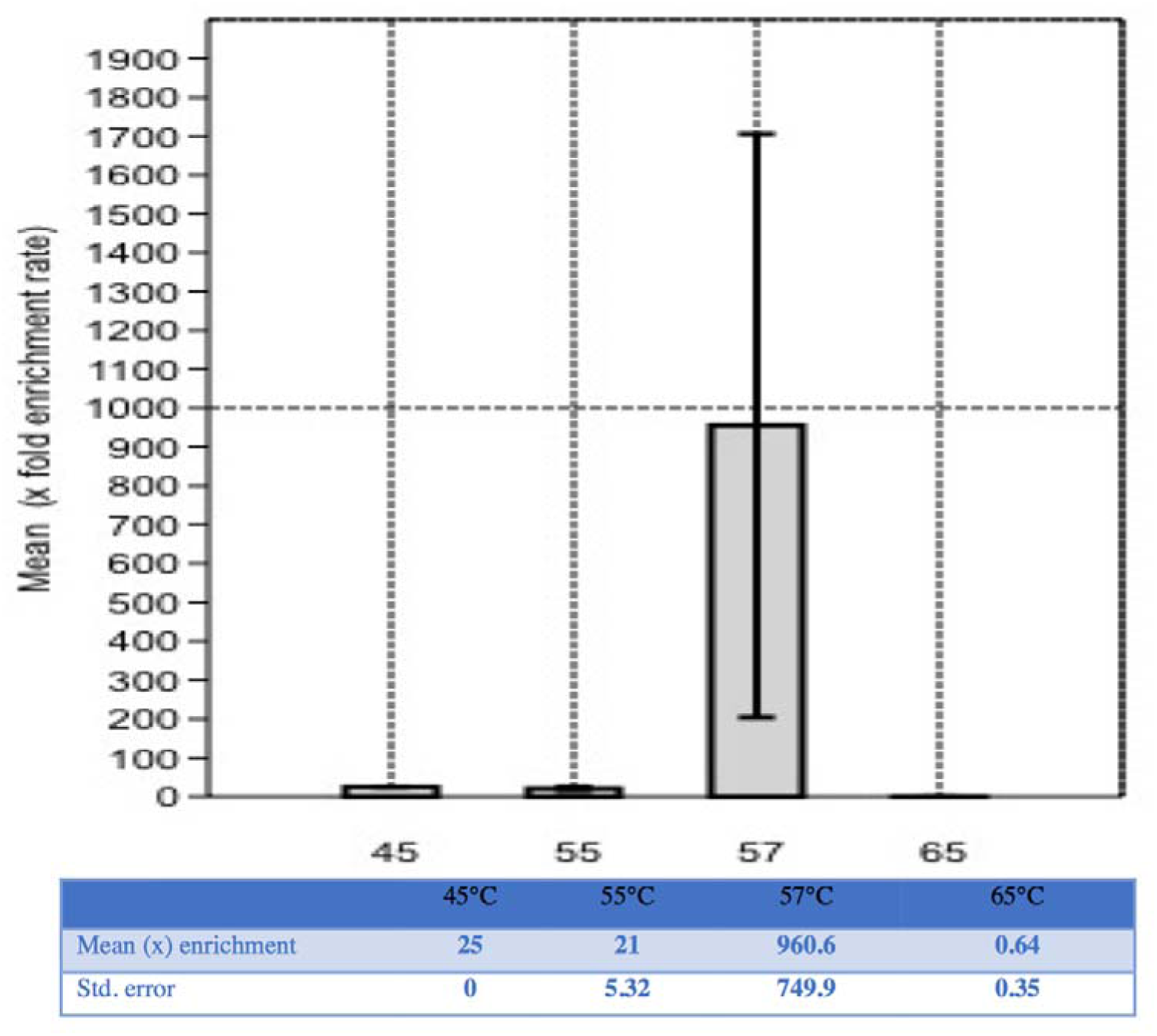
Optimisation of enrichment for ancient Sacred Ibis mitochondrial DNA. All libraries were constructed using a NEBNext kit with modifications by Meyer and Kircher (2010). Mean fold enrichment with Std. error are shown for ancient Sacred Ibis DNA hybridised using the following conditions: A) Hybridisation at 45°C; amplified using Phusion polymerase; Sequencing performed on HiSeq2000; B) Hybridisation at 55°C; amplified using Phusion polymerase; Sequencing performed on HiSeq2000; C) Hybridisation at 57°C; amplified using KAPA HiFi polymerase; Sequencing performed on MiSeq; D) Hybridisation at 65°C; amplified using Phusion polymerase; Sequencing performed on MiSeq.

### Clonality After Capture

The percentages of clonal sequences (not unique) were calculated for the ancient samples from the total mapped reads. We found that with ancient samples, post-capture libraries had higher clonality than pre-capture libraries (Fig. 4). We also found the lower the endogenous content of the ancient pre-capture libraries, the higher the post-capture clonality. This increase in clonality percentage is almost certainly due to the loss of sequences during the washes, as more amplification cycles were required post-capture.

**Fig. 4:**
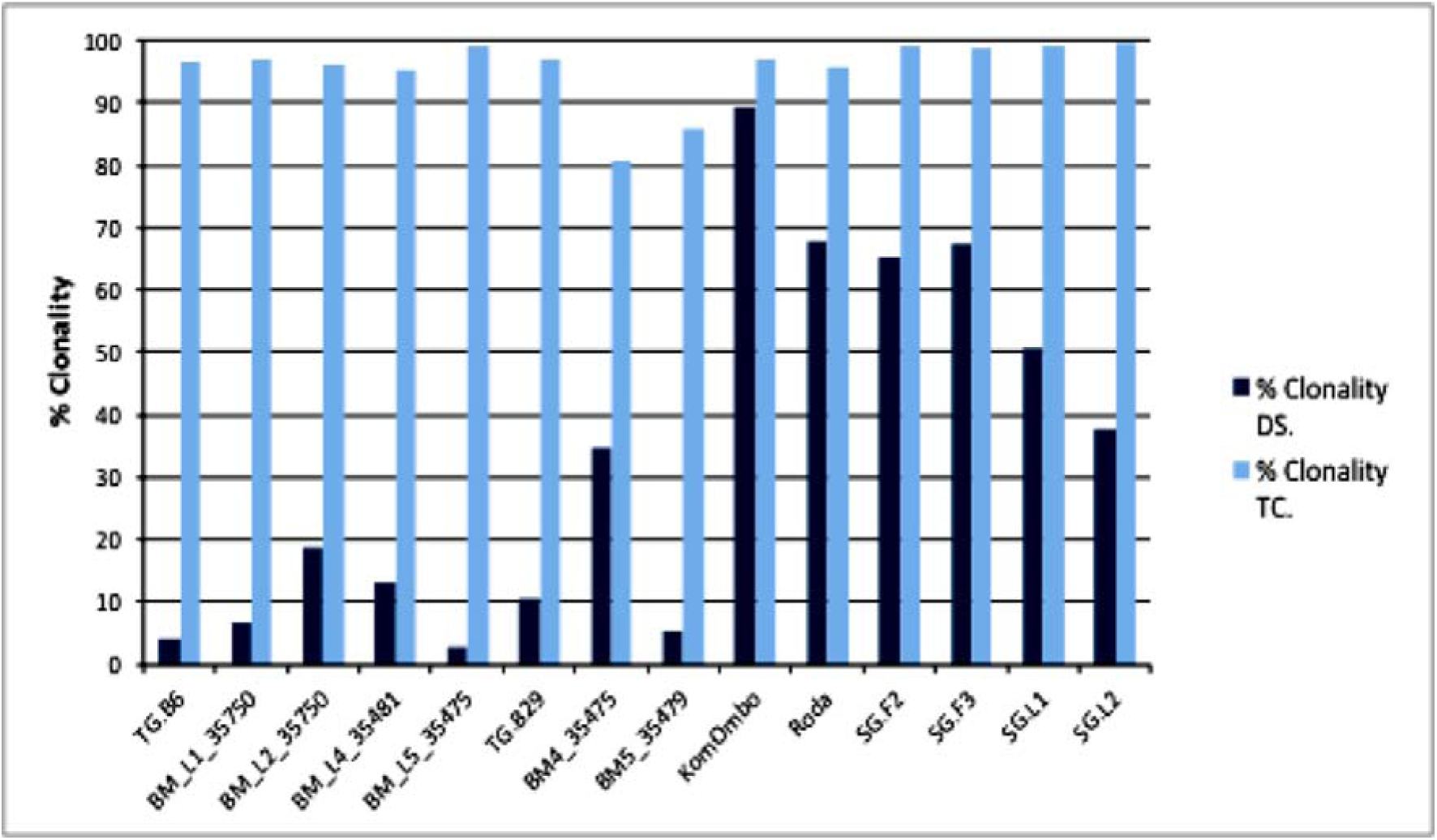
Percentage Clonality for Ancient Sacred Ibis libraries. The bars represent the % clonality for pre-capture shotgun sequenced libraries (DS) vs. Percentage Clonality after target capture for the same libraries (TC).

## Conclusion

The feasibility of retrieving authentic and informative DNA sequences from ancient materials originating from Egypt has been debated (Gilbert et al. 2005). We show that it is possible to retrieve significant amounts of endogenous DNA from ancient Sacred Ibis mummified material from deep within Egyptian catacombs. This is particularly surprising because Egyptian animal mummies in general were not mummified with the same care and attention shown towards royal or other human mummies.

We show that recent advances in ancient DNA extraction and library construction can result in the successful retrieval of ancient DNA from a number of Egyptian Ibis tissues such as bone, feather, tissue, and toe pad. Our results show that it was possible to achieve up to 4705-fold enrichment of highly contaminated and fragmented Egyptian ancient DNA using a combination of parameters, including the use of a modified DNA extraction method (Dabney, et al. 2013), an efficient ancient DNA library building protocol (Meyer & Kircher 2010), the polymerase KAPA HiFi Uracil+, and a hybridisation temperature of 57°C.

Sacred Ibis mitochondrial DNA was enriched using the MYbaits enrichment protocol. We examined the significance of temperature as a main, but not the sole, parameter to maximise the recovery of endogenous DNA from Sacred Ibis mummies. Our results show that the recovery of unique target sequences of ancient samples was significantly influenced by minor changes of the hybridisation conditions, as well as both the after-capture wash temperature used. Post capture temperature was always kept the same as the hybridisation temperature in use to maintain a higher stringency when washing away the contaminant sequences. For ancient Egyptian samples, hybridisation at 57°C, lower than the recommended 65°C, appears ideal. This lower temperature allowed more on-target specificity for fragmented and damaged sequences. However, reducing the hybridisation temperature to 45°C, leads to the loss of selectivity of baits in the case of the Sacred Ibis mitogenome. This illustrates that decreased hybridisation temperatures are beneficial for ancient samples.

## Acknowledgement

We are grateful to Human Frontier Science for financial support in the form of grant to PIs, Lambert, Ikram, Holland, and Willerslev. We are also grateful to The Danish National High-Throughput DNA Sequencing Centre for sequencing the samples. We would like to thank Caitlin Curtis for her support. SW is really appreciative to the Ministry of Antiquities for permitting ancient samples collection from the catacombs; also to Al Kasr Al Ani Medical School for allowing her to use their ancient DNA laboratory. A number of museums kindly provided material for this study including: The British Museum, The Ancient Egyptian Animal Bio Bank at Manchester Museum, Manchester, UK, especially Dr. Lidija M. McKnight and the Musée des Confluences, Lyon, France, particularly Stéphanie Porcier. SW thanks Griffith University for a PhD scholarship and DML and SW thank the Environmental Futures Research Institute and Griffith University for additional support.

